# Diversity patterns of marine heterotrophic culturable bacteria along vertical and latitudinal gradients

**DOI:** 10.1101/774992

**Authors:** Isabel Sanz-Sáez, Guillem Salazar, Elena Lara, Marta Royo-Llonch, Dolors Vaqué, Carlos M. Duarte, Josep M. Gasol, Carlos Pedrós-Alió, Olga Sánchez, Silvia G. Acinas

**Author notes:** Correspondence: Olga Sánchez; Silvia G. Acinas.

## Abstract

Nowadays, there is a significant gap in the knowledge of the diversity and patterns for marine heterotrophic culturable microorganisms. In addition, most of the bacterial isolation efforts have focused on the photic ocean leaving the deep ocean less explored. We have isolated 1561 bacterial strains covering both photic (817) and aphotic layers (744) including isolates from the oxygen minimum zone (362) and the bathypelagic (382) from a variety of oceanographic regions including the North Western Mediterranean Sea, the North and South Atlantic Oceans, the Indian, the Pacific, and the Arctic Oceans. The partial sequencing of the 16S rRNA gene of all isolates revealed that they mainly affiliate with the classes *Alphaproteobacteria* (35.9%) and *Gammaproteobacteria* (38.6%), as well as, phylum *Bacteroidetes* (16.5%). The genera *Alteromonas* and *Erythrobacter* were the most widespread heterotrophic bacteria in the ocean able to grow on solid agar media. When comparing the sequences of all isolates, 37% of them were 100% identical. In fact, we found that 59% of the total aphotic isolates were 100% identical to photic isolates, indicating the ubiquity of some bacterial isolates along the water column. Unweighted UniFrac distances did not show significant differences among stations regardless of their geographic distance or depth, reflecting the wide dispersion of the culturable bacterial assemblage. This isolates collection provides an overview of the distribution patterns of cosmopolitan marine culturable heterotrophic bacteria.

## Introduction

High-throughput sequencing (HTS) studies of ribosomal genes [1] have provided great advances in the knowledge of microbial diversity in multiple environments including marine ecosystems [2–4]. They allowed to study the richness and community composition of marine bacterioplankton communities, as well as, the identification and functional insights of the most abundant microorganisms [5–7]. Different main patterns emerged from 16S rRNA amplicon tags (16S iTAGs) and metagenomic (16S miTAGs) studies applied to ocean ecosystems: (i) the presence of few abundant operational taxonomic units (OTUs) and a long tail of rare and low abundant OTUs, called the “rare biosphere” [1, 8, 9], (ii) an increase of richness in deeper layers [3, 10], (iii) the vertical segregation between photic and aphotic samples [3] but also vertical connectivity between surface and deep ocean taxa through sinking particles [11], (iv) the limited dispersion of bacterial communities across space [3, 12, 13] and, (v) the presence of few cosmopolitan taxa [14, 15]. However, there is still a significant gap in the knowledge of the diversity of culturable microorganisms in the ocean, and if similar patterns are applied to this fraction of the bacterioplankton community, specifically covering broader gradients. Also, it is unclear which are the most cosmopolitan culturable marine heterotrophic bacteria in the ocean.

On top of that, most of the studies targeting the marine heterotrophic culturable bacteria have focused on the upper ocean (0-200 m depth) or on specific oceanographic regions [16–19], while studies covering vertical gradients are less frequent [20–22]. Efforts to culture bacteria from the deep ocean (>200 m) have focused mostly on isolates from hydrothermal vents [23–25], whale carcasses [26], trenches [27], and deep-sea sediments [22, 28–31]. Thus, very few studies have analyzed the diversity of isolates from mesopelagic (in particular the oxygen minimum zone (OMZ) [32–34]), the bathypelagic and abyssopelagic waters [20, 35–37], and those available were mainly done at a local or regional scale. Thus, a study of the culturable microorganisms covering vertical gradients including underexplored areas such as the OMZ and the bathypelagic areas is missing.

Here we present an extensive marine heterotrophic bacterial culture collection with 1561 marine bacteria retrieved from different seawater depths covering also a large geographical extent and latitude. Through the analysis of their partial 16S rRNA sequences (average 526 bp), we aim to describe some general patters of isolated bacteria rather than describe new bacterial species. To do that we used a well stablished marine solid medium in order to describe that fraction of the bacterioplankton community than can be commonly isolated under laboratory conditions (nutrient rich medium, standard oxygen concentrations and atmospheric pressure). We have revealed both the most cosmopolitan and the locally distributed heterotrophic culturable bacteria across oceans, and revealed some different patterns of vertical distribution showing large connectivity and dispersion of the isolated heterotrophic fraction.

## Materials and methods

### Study areas and sampling

A total of eight photic-layer, four OMZ, and seven bathypelagic samples were taken during different oceanographic cruises in several sampling stations distributed along a wide range of latitudes (Fig. 1). Photic layer samples (Table 1) were collected in the Atlantic and Indian Oceans during the *Tara* Oceans expedition in 2009-2013 [38], and from the Arctic Ocean during the ATP 09 cruise in 2009 [39]. Additionally, surface seawater samples from the Blanes Bay Microbial Observatory (BBMO, http://www.icm.csic.es/bio/projects/icmicrobis/bbmo) in the NW Mediterranean Sea were collected in May 2015. Oxygen minimum zone samples (Table 1) were taken from the Indian and Pacific Oceans also during the *Tara* Oceans expedition in 2009-2013 [38]. Bathypelagic samples (Table 1) from the Atlantic Ocean at ~ 4000 m depth were taken from six different stations during the Malaspina 2010 Circumnavigation Expedition [40]. One of the stations sampled was located in the North Atlantic Ocean, whereas the other five stations were located in the South Atlantic Ocean; one of them was particularly placed in the Agulhas Ring, where deep waters from the South Atlantic converge and mix with Indian Ocean deep-water masses [41]. In addition, one bathypelagic sample was collected at 2000 m depth in the NW Mediterranean during the MIFASOL cruise in September 2014.

**Table 1.** Characteristics of the different samples used for isolation of bacteria. Non-redundant isolates stand for those isolates in the same sample that presented 100% similarity in their partial 16S rRNA sequences.

| Cruise | Station | Oceanic location | Latitude | Longitude | Depth (m) | *In situ*<br>temperature<br>(°C) | Nº of<br>sequenced<br>isolates | Nº of non-redundant<br>isolates |
| --- | --- | --- | --- | --- | --- | --- | --- | --- |
| Tara Oceans | ST 39 | Indian Ocean | 19° 2.24' N | 64° 29.24' E | 5.5 | 26.2 | 104 | 25 |
|  | ST 39 | Indian Ocean | 18° 35.2' N | 66° 28.22' E | 25 | 26.8 | 243 | 53 |
|  | ST 39 | Indian Ocean | 18° 43.12' N | 66° 21.3' E | 268.2 | 15.6 | 88 | 18 |
|  | ST 67 | South Atlantic | 32° 17.31' S | 17° 12.22' E | 5 | 12.8 | 115 | 49 |
|  | ST 72 | South Atlantic | 8° 46.44' S | 17° 54.36' W | 5 | 25 | 71 | 33 |
|  | ST 76 | South Atlantic | 20° 56.7' S | 35° 10.49' W | 5 | 23.3 | 89 | 27 |
|  | ST 151 | North Atlantic | 36° 10.17' N | 29° 1.23' W | 5 | 17.3 | 76 | 33 |
|  | ST 102 | Pacific Ocean | 5° 16.12' S | 85° 13.12' O | 475.6 | 9.2 | 97 | 15 |
|  | ST 111 | Pacific Ocean | 16° 57.36' S | 100° 39.36' O | 347.1 | 10.9 | 98 | 35 |
|  | ST 138 | Pacific Ocean | 6° 22.12' N | 103° 4.12' O | 444.9 | 8.2 | 79 | 34 |
| ATP | AR_1 | Arctic Ocean | 78° 20.00' N | 15° 00.00' E | 2 | 6.2 | 13 | 9 |
|  | AR_2 | Arctic Ocean | 76° 28.65' N | 28° 00.62' E | 25 | -1.2 | 20 | 9 |
| Malaspina | ST 10 | North Atlantic | 21° 33.36' N | 23°26' W | 4002 | 2.04 | 20 | 9 |
|  | ST 17 | South Atlantic | 3° 1.48' S | 27° 19.48' W | 4002 | 1.74 | 93 | 24 |
|  | ST 23 | South Atlantic | 15° 49.48' S | 33° 24.36' W | 4003 | 1.45 | 94 | 39 |
|  | ST 32 | South Atlantic | 26° 56.8' S | 21° 24' W | 3200 | 2.5 | 39 | 16 |
|  | ST 33 | South Atlantic | 27° 33.2' S | 18° 5.4' W | 3904 | 1.7 | 5 | 5 |
|  | ST 43 | South Atlantic | 32° 48.8' S | 12° 46.2' E | 4000 | 1.2 | 4 | 4 |
| Mifasol | ST 8 | NW Mediterranean | 40° 38.41' N | 2° 50' E | 2000 | 13.2 | 127 | 36 |
| BBMO | IBSURF | NW Mediterranean | 41° 40' N | 2° 48' E | 5 | 17.71 | 86 | 43 |

**Fig. 1.**
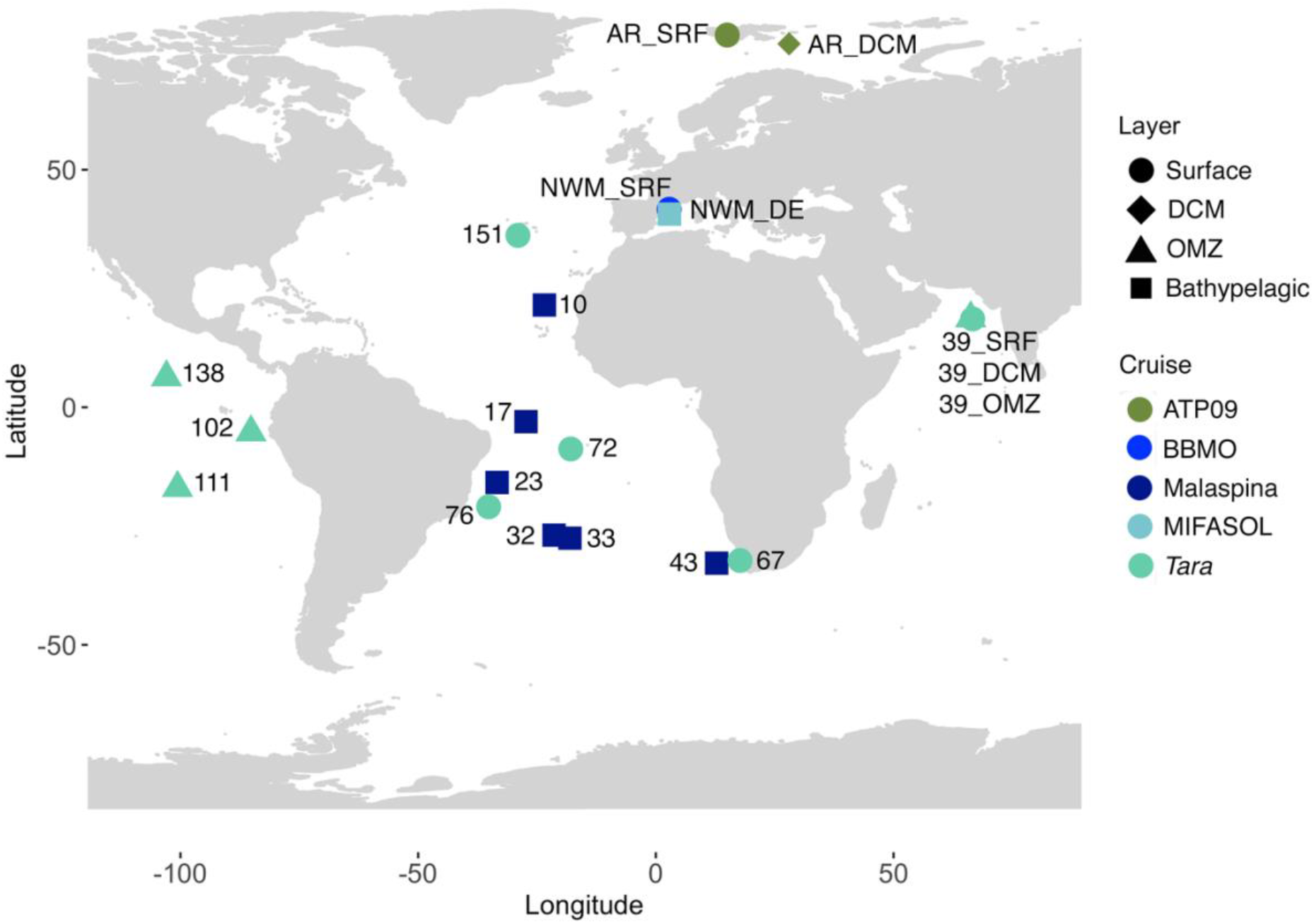
Map showing the sampling stations of the present study. Circles indicate, surface stations; diamonds, deep chlorophyll maximum (DCM) stations; triangles, oxygen minimum zone (OMZ) stations; and squares, bathypelagic stations. The different colours indicate the oceanographic cruise in which the samples were collected: dark green, ATP09; blue, Blanes Bay Microbial Observatory (BBMO); dark blue, Malaspina; light blue, MIFASOL; and turquoise, *Tara* Oceans.

In each of these stations, seawater was collected using Niskin bottles attached to a rosette sampling system, except at BBMO, where samples were collected with a bucket. Seawater was sequentially filtered through 200 μm and 20 μm meshes to remove large plankton cells. Two subsamples were kept in 2 ml Eppendorf tubes with DMSO 7% final concentration and stored at −80ºC until further processing in the laboratory.

Geographical coordinates of stations, sampled depth, *in situ* temperature and total number of sequenced isolates are listed in Table 1.

### Culturing and isolation

Isolates were obtained by plating 100 μl of undiluted and 10x diluted seawater from the photic, OMZ and bathypelagic samples, in triplicates, onto Zobell agar plates (i.e. 5g peptone, 1g yeast extract and 15 g agar in 750 ml of 30 kDa filtered seawater and 250 ml of Milli-Q water) or Marine Agar 2216 plates, which is based also on the Zobell medium formulation [42] (5 g peptone, 1 g yeast extract, 0.1 g ferric citrate, 19.4 g sodium chloride, 8.8 g magnesium chloride, 3.24 g sodium sulphate, 1.8 g calcium chloride, 0.5 g potassium chloride, 0.16 g sodium bicarbonate, 0.08 g potassium bromide, 0.034 g strontium chloride, 0.022 g boric acid, 0.004 g sodium silicate, 0.0016 g ammonium nitrate, 0.008 g disodium phosphate, 0.0024 g sodium fluorate, 15 g agar in 1 L of distilled water). Our medium culturing strategy was only focused to retrieve heterotrophic marine bacteria that could grow easily under laboratory conditions (nutrient rich media, standard oxygen concentrations and atmospheric pressure) using two similar culturing media. The only difference between Zobell agar and Marine Agar 2216 plates is the use of natural seawater (Zobell agar), or the addition of the minerals and salts contained in natural seawater to distilled water (Marine Agar 2216). Indeed, we did not observe significant differences (Fisher test analyses, data not shown) in the bacterial isolation between both media.

Photic layer and OMZ samples were incubated at room temperature (RT), while bathypelagic samples were incubated at their *in situ* temperature, which ranged from ~4ºC in the (Atlantic Ocean at 4000 m depth) to 12ºC (NW Mediterranean at 2000 m depth) (Tables 1 and S1), but also at RT in order to assure bacterial recovery from all stations. In all cases, triplicates of each temperature condition and dilution were incubated in the dark until no more colonies appeared (10-30 days).

A total of 1561 bacterial isolates were randomly selected for DNA amplification and partial sequencing of their 16S rRNA gene (Table 1 and details below). Similar number of isolates were sequenced from photic layers (817; average: 102 isolates per station) and from deep oceans (744; average: 67 isolates per station) with 362 isolates from the OMZ and 382 from the bathypelagic. In most of the bathypelagic samples we collected all colonies appearing in the plates, which ranged from 6 to 129. Colonies were streaked on agar plates in duplicate to ensure their purity and avoid contamination. The isolates were stored in the broth medium used with glycerol 25% in cryovials at −80ºC.

### PCR amplification and sequencing of the 16S rRNA gene

Available DNA used for template in Polymerase Chain Reaction (PCR) was extracted from 200 μL of isolates liquid cultures placed in 96 well plates, diluted 1:4 and heated (95ºC, 15 min) to cause cell lyses. The partial 16S rRNA gene sequences were PCR amplified using bacterial primers 358F (5’-CCT ACG GGA GGC AGC AG-3’) [43] and 907Rmod (5’-CCG TCA ATT CMT TTG AGT TT-3’) [44]. The complete 16S rRNA gene was amplified in specific strains after DNA extraction using the DNeasy Blood & Tissue kit (Qiagen), following the manufacturer’s recommendations, and using the modified primers from Page et al. [45] 27F (5’-AGR GTT TGA TCM TGG CTC AG −3’) and 1492R (5’-TAC GGY TAC CTT GTT AYG ACT T −3’). Each PCR reaction with a final volume of 25 μl contained: 2 μl of template DNA, 0.5 μl of each deoxynucleotide triphosphate at a concentration of 10 μM, 0.75 μl of MgCl21.5 mM, 0.5 μl of each primer at a concentration of 10 μM, 0.125 μl of Taq DNA polymerase (Invitrogen), 2.5 μl of PCR buffer supplied by the manufacturer (Invitrogen, Paisley, UK) and Milli-Q water up to the final volume. Reactions were carried out in a Biorad thermocycler using the following program: initial denaturation at 94ºC for 5 min, followed by 30 cycles of 1 min at 94ºC, 1 min at 55ºC and 2 min at 72ºC, and a final extension step of 10 min at 72ºC. The PCR products were verified and quantified by agarose gel electrophoresis with a standard low DNA mass ladder (Invitrogen). Purification and OneShot Sanger sequencing of 16S rRNA gene products was performed by Genoscreen (Lille, France) with primer 358F for partial sequences, and with both 27F and 1492R for complete sequences. ChromasPro 2.1.8 software (Technelysium) was used for manual cleaning and quality control of the sequences.

### Data processing and taxonomic classification

The 16S rRNA sequences of our cultured strains were clustered at 99% sequence similarity [46] in order to define different operational taxonomic units (OTUs or isolated OTUs) and construct OTU-abundance tables for the different stations and layers studied (Table S2) using UCLUST algorithm from the USEARCH software [47]. The different OTUs were taxonomically classified using the least common ancestor (LCA) method in SINA classifier [48], using both Silva (release 132 in 2017) and RDP (Ribosomal Database Project, release 11) databases. Parallelly, isolates sequences were submitted to BLASTn [49] with two subsets of the RDP database. One including only the uncultured bacteria (Closest Environmental Match, CEM), and another including only the cultured bacteria (Closest Cultured Match, CCM) in order to extract the percentages of similarity with both datasets (Table S3 and Table S4), and to assess whether our isolates were similar to effectively published cultured organisms.

Additionally, a more restrictive clustering at 100% sequence similarity (USEARCH software) was used to define OTUs and to detect how many bathypelagic, OMZ and photic-layer bacterial isolates were identical, and thus, to identify bacterial taxa or strains that could distribute along all the water column. Such comparisons were done with: (i) photic and bathypelagic isolates sequences retrieved from neighbouring stations and (ii) the whole isolates dataset.

### Comparisons between layers and statistical analyses

To infer differences between the diversity and richness of the isolates obtained along the oceanic vertical gradient, the OTU-abundance table constructed with the sequences clustering at 99% was sampled down to the lowest sampling effort (362 isolates in the OMZ). In this manner, the rarefied or subsampled OTU table was obtained using the function *rrarefy.perm* with 1000 permutations from the R package *EcolUtils* [50], while rarefaction curves to estimate the sampling effort were performed with the package *vegan* 2.5-3 [51]. Calculation of Chao1 and ACE species richness estimates, and Shannon-Weaver and Simpson diversity index estimates were also carried out with the rarefied OTU table using the R package *vegan* version 2.5-3.

The coverage (C) for each of the depths was also calculated by the equation 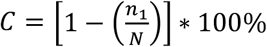, where *N* is the number of OTUs being examined and *n1* represents the number of OTUs occurring only once or singletons [52].

Fisher tests were performed in R package *stats* version 3.4.3 [53] and were used to: (i) detect taxa with a significant presence at a specific depth in those truly comparable stations, that is stations which present photic and bathypelagic samples from neighbouring stations, and (ii) evaluate this significant presence of certain taxa in a specific layer including all photic, OMZ and bathypelagic samples. *P*-values of the Fisher test were adjusted for multiple comparisons using the Hochberg False Discovery Rate correction (FDR). To assess significance, the statistical analyses were set to a conservative alpha value of 0.01.

### Phylogenetic analysis

In order to see phylogenetic relationships and explore the connectivity between photic-layer, OMZ and bathypelagic isolates, as well as, to detect possible novel strains, phylogenetic trees for *Alphaproteobacteria*, *Gammaproteobacteria*, *Bacteroidetes* and Gram positive bacteria (Fig. S1a-d) were built. They included the non-redundant sequences dataset of each station and the taxonomic affiliation of their Closest Cultured Match (CCM) and Closest Environmental Match (CEM) obtained after BLASTn search against the RDP databases (Table S3 and S4). The total pool of sequences were first aligned with the SINA web alignment tool (http://www.arb-silva.de/aligner/) [48] and imported into the phylogenetic software MEGA 5.2.2 [54]. The phylogenetic trees were constructed with the Neighbour Joining (NJ) algorithm using the Jukes-Cantor distance and 1000 bootstrap replicates.

Additionally, to support the novelty of putative new species or genera (isolates that presented a percentage of similarity below the 97% with public databases), the complete 16S rRNA gene was sequenced for one isolate, with which two more strains clustered at 100% similarity, and phylogenetic trees were constructed as described above. The trees included the complete and partial sequences, the best hits from uncultured and cultured microorganisms, extracted from local alignments against RDP 11, Silva LTP (Living Tree project), and National Center for Biotechnology Information (NCBI) databases, and the reference 16S rRNA genes from their related genera.

### Analyses of the genetic distances between retrieved isolates

Unweighted UniFrac distances between isolates of the different stations were calculated using the R version 3.4.3 [53] package *picante* [55]. The analysis required a phylogenetic tree including all the isolates sequences, and an outgroup sequence (*Rhodopirellula baltica*, accession number EF012748), which was constructed as mentioned above, and an OTU-abundance table constructed at 100% sequence similarity using the UCLUST algorithm from the USEARCH software [47] indicating the presence of the different OTUs in each of the samples studied. The resulting UniFrac distance matrix was used to estimate genetic distances between samples from different depths and to perform a non-metric multidimensional scaling analysis [56] using the R package *vegan* 2.5-3 [51]. One-way analysis of variance (ANOVA) test followed by the post hoc Tukey’s Honestly significant difference (HSD) test from the R package *stats* version 3.4.3 [53] was perform to see significative differences between groups. To asses significance, the statistical analyses were set to a conservative alpha value of 0.05. Furthermore, we performed different analyses to test whether environmental parameters (temperature, salinity) and Chlorophyll a (*Chl* a) could explain the patterns of genetic distance between samples of different depths: (i) a Mantel test correlation with the R package *ape* version 5.1 [57], including the unweighted UniFrac distance matrix and a distance matrix based on Euclidian distances between the scaled environmental parameters, and (ii) a distance-based redundancy analysis (dbRDA, [58]) with the R package *vegan* version 2.5-3.

### Nucleotide sequences accession number

The 16S rRNA gene sequences of the bacterial isolates retrieved in this study were deposited in GenBank. Sequences from all isolates, except those coming from the OMZ regions and those from the surface Indian Ocean, are deposited under the following accession numbers MH731309 - MH732621. Notice that among these accession numbers other isolates are included, originated from the same locations but isolated with another culture medium and not included in this study. Isolates retrieved from the OMZ and those from the surface Indian Ocean are deposited under the following accession numbers MK658870-MK659428.

## Results

### Bacterial richness and diversity patterns of heterotrophic isolates

The partial 16S rRNA sequences of the cultured strains were grouped into operational taxonomic units (isolated OTUs, referred hereafter as OTUs) using similarity thresholds at 99% (OTUs99) and 100% (OTUs100). Those OTUs were then compared and analysed.

The number of OTUs detected and the observed richness (S.obs) was very similar in all layers for both OTUs100 and OTUs99, being the OMZ the layer with the lowest observed values (Table 2). Rarefaction curves for the non-rarefied (Fig. 2a) and rarefied (Fig. 2b) OTU tables showed slightly higher richness for the photic samples compared to the OMZ and bathypelagic but they did not reach an asymptote. Thus, we calculated the species richness estimates (Chao-1, ACE) and they increased with depth for OTUs100 (from 140 in the photic up to 174 in the bathypelagic), but it was in the OMZ where we found higher values for the OTUs99 (Table 2). Nevertheless, Shannon-Weaver and Simpson diversity indices showed, for both clustering thresholds, a slightly higher diversity for photic layers. Good’s coverage analyses per layer ranged from 48.8 to 55.8% (OTUs100) and from 56.1 to 70.5% (OTUs99)(Table 2). These results indicated that the isolates dataset, even if not saturated, represents a reasonable inventory of the heterotrophic marine bacteria.

**Table 2.** Richness, Diversity and Coverage of the isolates retrieved per depth. Results of diversity indices calculated with the OTU-abundance tables per layer constructed at 99% and 100% sequences similarity and rarefied down to the lowest isolated layer (OMZ with 362 isolates). OMZ: oxygen minimum zone; Singletons: OTUs appearing only once and S.obs: Observed species.

|  | 99% |  |  | 100% |  |  |
| --- | --- | --- | --- | --- | --- | --- |
|  | Photic | OMZ | Bathypelagic | Photic | OMZ | Bathypelagic |
| *Number of isolates* | 346 | 362 | 368 | 320 | 362 | 372 |
| *Number of OTUs* | 61 | 57 | 59 | 95 | 80 | 98 |
| *Number of Singletons* | 18 | 25 | 20 | 42 | 41 | 50 |
| *Coverage* | 70.5% | 56.1% | 66.1% | 55.8% | 48.75% | 49% |
| *S.obs* | 61 | 57 | 59 | 95 | 80 | 98 |
| *Chao1* | 71.2 | 99.9 | 74.8 | 140.31 | 162 | 174.6 |
| *ACE* | 77.8 | 87.6 | 81.9 | 145.56 | 143.3 | 179.9 |
| *Shannon-Weaver* | 3.4 | 3.15 | 3.3 | 4 | 3.5 | 3.87 |
| *Simpson* | 0.95 | 0.92 | 0.94 | 0.97 | 0.94 | 0.96 |

**Fig. 2.**
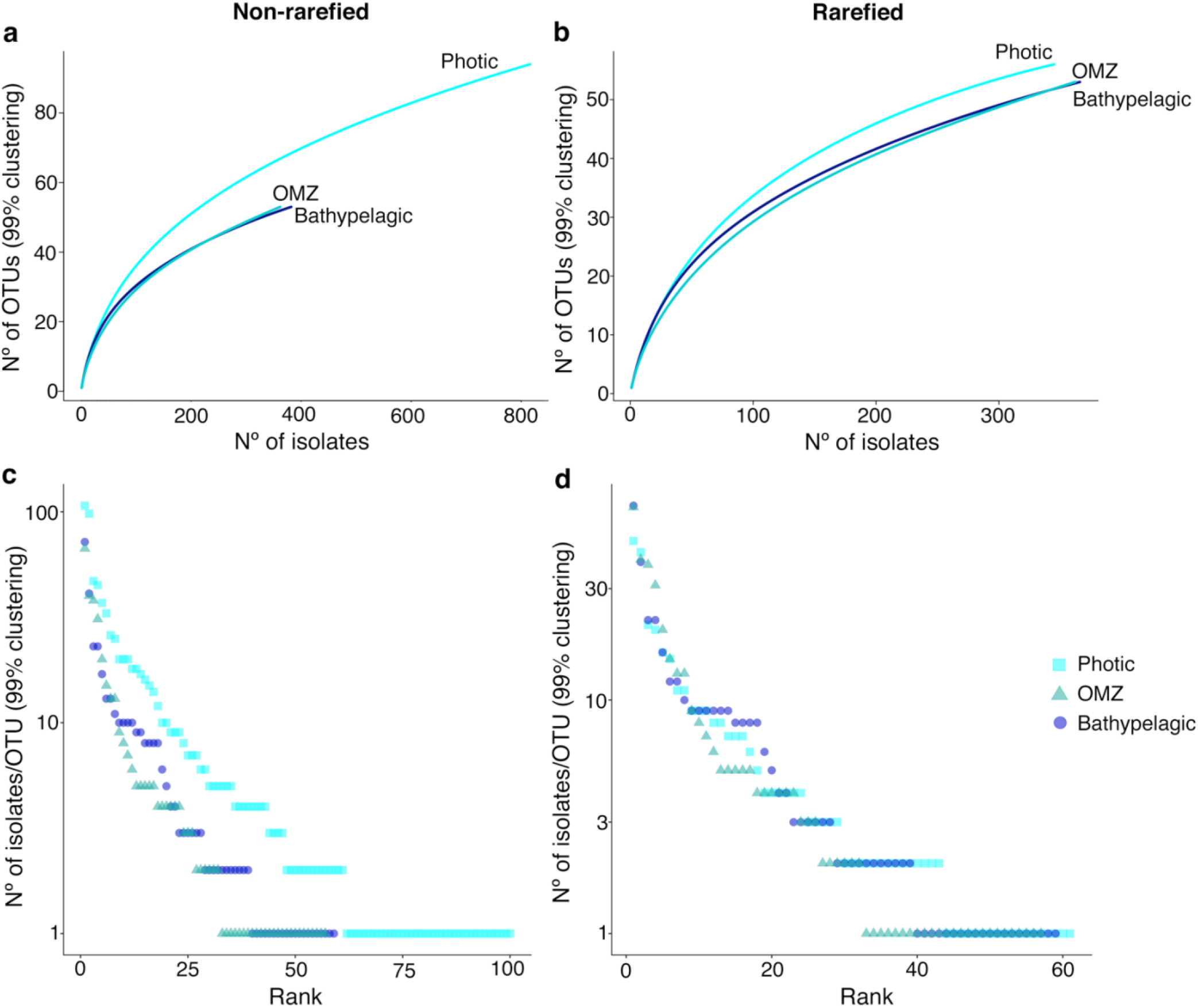
**(a-b)** Rarefaction curves for photic, OMZ and bathypelagic samples extracted also from the non-rarefied and rarefied OTU table. **(c-d)** Rank abundance plots showing the number of isolates per OTU (at 99% clustering) obtained in the three layers studied for non-rarefied OTU table, and rarefied OTU table down to the lowest isolated layer, in this case, the OMZ with 362 isolates. Y axis are in log10 scale. Colour and shape indicate the layer where the OTUs were isolated: Photic-layer, light-blue squares; Oxygen minimum zone (OMZ), Turquoise triangles; and Bathypelagic ocean, dark blue circles.

Finally, rank abundance plots of the non-rarefied (Fig. 2c) and rarefied (Fig. 2d) OTU tables presented, for the three depths studied, a steep curve, which is indicative of low evenness. Thus, there were a few abundant OTUs with a large number of strains and a large proportion of OTUs that had few representatives (rare OTUs).

### Shared diversity between photic and bathypelagic samples from neighbouring stations

The 382 16S rRNA sequences from the bathypelagic samples were compared to the 437 photic-layer sequences (excluding those from the Indian and Arctic Ocean because bathypelagic isolates could not be retrieved in these cases). The data were normalized down to the lowest isolated sequences that corresponded to the strains retrieved from the bathypelagic ocean, but very similar results were obtained with the rarefied and non-rarefied data (Table S5). Surprisingly, a large proportion (58.9%) of the bathypelagic sequences were 100% identical to those of the photic isolates. Further analyses in both sets of sequences were done at both OTU and genus taxonomic level (Fig. 3) and showed that photic isolates clustered in 73 different OTUs while bathypelagic isolates clustered in 56 OTUs (99% sequence similarity). Slightly higher numbers of *Gammaproteobacteria*, *Alphaproteobacteria*, and *Bacteroidetes* OTUs were retrieved from the photic layer (Fig. 3a). More differences were found in the *Actinobacteria*, dominated by only photic-layer OTUs, and *Firmicutes*, present only in the bathypelagic. Furthermore, when we analyzed together the photic and bathypelagic isolates and we defined OTUs at 99% sequence similarity, we found 29 OTUs which included strains of both layers (Fig. 3b). These shared OTUs included 68.9% of the sequences (Fig. 3b) and an average number of 18.2 sequences per OTU. Despite the fact that this large group of OTUs was found in both layers, 41 OTUs, representing 18.3% of the sequences, were still found only in the photic zone, with an average number of 3.4 sequences per OTU. On the other hand, 28 OTUs (12.8% of the sequences) were found only in the bathypelagic ocean, with an average of 3.5 sequences per OTU (Fig. 3b)(Table S6). At the genus level we found 23 genera common to both layers (including 80.4% of the analysed sequences), 10 (4.6%) only in the bathypelagic zone and 24 (15%) among the photic isolates (Table S7). Some of the isolates that were only recovered from the bathypelagic waters belonged to *Bacillus*, *Loktanella, Novosphingobium*, *or Roseovarius*, whereas all the genera affiliating with the *Actinobacteria* phylum and other genera, including *Maricaulis*, *Polaribacter*, *Roseivirga, Vibrio* or *Winogradskyella*, were only isolated from the photic-layer within our dataset. Nevertheless, Fisher tests (Table S8) indicated a significative presence of *Roseivirga*, *Winogradskyella*, and *Vibrio* in the photic layer, while *Sulfitobacter*, despite being found in both layers, was more prone to be isolated from the bathypelagic samples.

**Fig. 3.**
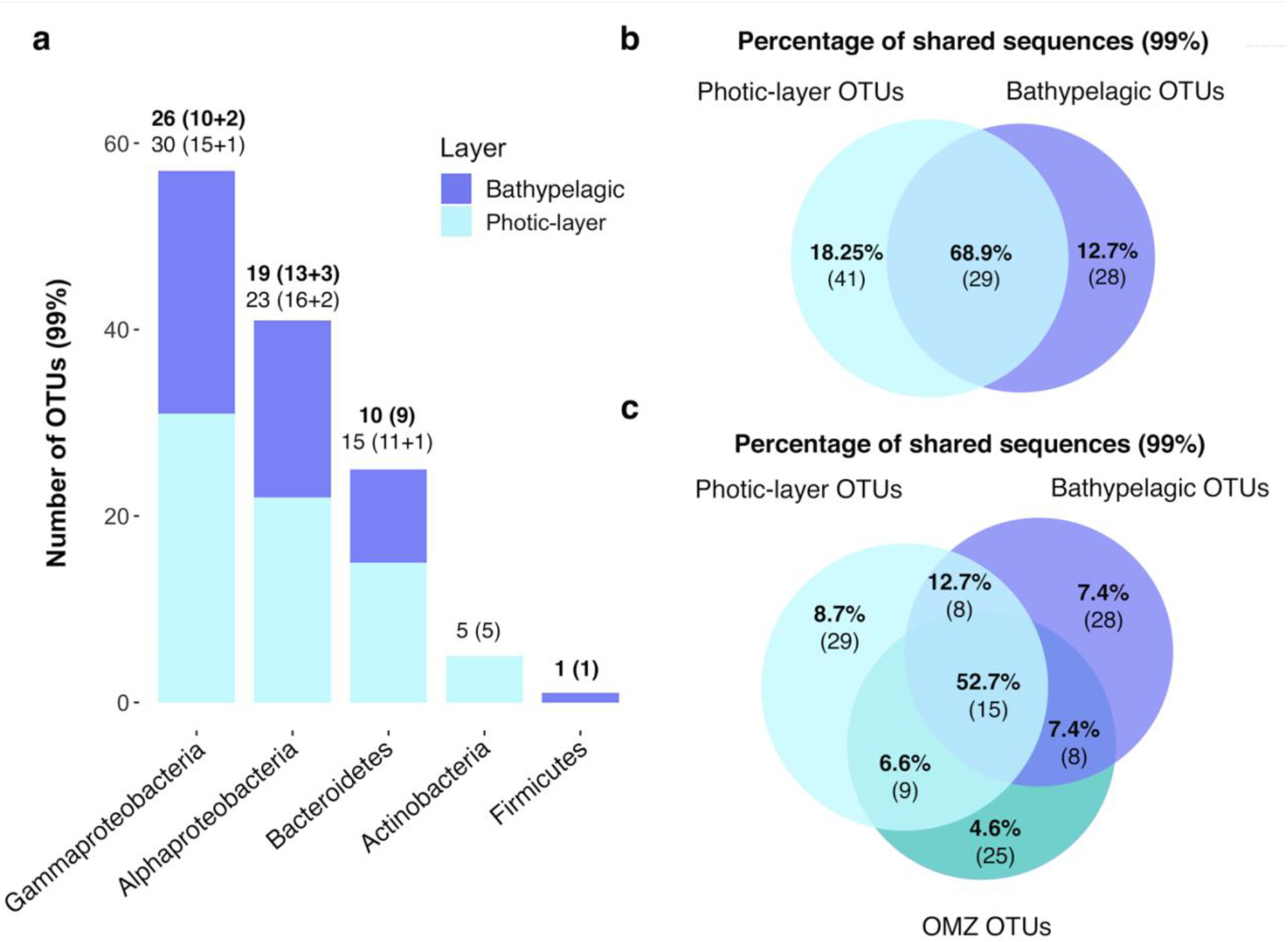
OTUs retrieved from photic-layer and deep-sea waters. **(a)** Phyla/Class retrieved per oceanic layer. Each bar of the plot represents a phylum, or class in the case of *Proteobacteria*. OTUs were defined by clustering at 99% sequence similarity. The dark blue represents the total number of OTUs retrieved from bathypelagic samples, whereas the light blue represents the total number of OTUs isolated from photic samples. The number of OTUs is shown above each bar. In bold, the ones from the bathypelagic, and in regular type, those recovered from the photic samples. Numbers between brackets indicate the number of genera that represent those OTUs plus the unclassified at the genus level. **(b)** Venn diagram indicating the percentages of the sequences shared between photic and bathypelagic samples from neighbouring stations. Numbers inside brackets indicate the number of shared OTUs corresponding to that percentage of sequences. **(c)** Venn diagram indicating the percentages of the sequences shared between photic, OMZ and bathypelagic layers. Numbers inside brackets indicate the number of shared OTUs corresponding to that percentage of sequences. Indicated numbers in all Venn diagrams are extracted from the rarefied OTU-abundance tables.

### Shared diversity between all photic, OMZ, and bathypelagic samples across oceans

We explored the similarity between OTUs from different layers (photic, OMZ and bathypelagic) and across distant oceans covering large spatial and latitudinal scales. The non-rarefied OTU table (OTUs99), including all the photic (817), OMZ (362), and bathypelagic (382) isolates, and the rarefied OTU table down to the lowest isolated depth (OMZ) were used for analyses, and because minor differences were observed (Table S9 and S10), the results mentioned here refer only to those obtained with the rarefaction. The 122 OTUs obtained, revealed that 15 OTUs (Fig. 3c) included isolates from all layers, accounting for 52.7% of the total sequences (Fig. 3c). Further, eight OTUs (12.7% of the sequences) were common to photic and bathypelagic isolates, nine (6.6%) to photic and OMZ isolates, and eight (7.36%) to OMZ and bathypelagic isolates (Fig. 3c). Nevertheless, a substantial proportion of isolates were only retrieved from one of the layers: 29 OTUs were only found in the photic samples, 25 in the OMZ, and 28 in the bathypelagic samples (Fig. 3c)(Table S11). The taxonomic classification of these OTUs, using the LCA method, designated a total of 59 different genera and 10 OTUs that could not be classified at the genus level. From these 59 genera, 13 genera were found along all the water column including 75% of the total isolates, and between four and six genera were shared by two layers. On the other hand, the photic ocean was again the layer with the highest number of retrieved genera that were not observed in the other two depths, even though they accounted for only 5.6% of the sequences (Table S12). Interestingly, Fisher tests applied to the genera retrieved indicated again more recovery of *Roseivirga*, *Vibrio* and *Winogradskyella* in the photic layers and of *Sulfitobacter* in the bathypelagic. Additionally, some other genera that were mostly isolated from only one of the depths within our dataset seemed to have also a significative presence in a specific layer, such as *Nereida*, *Maricaulis* (Photic), *Gramella* (OMZ), or *Joostella* (Bathypelagic) (Table S13).

### Cosmopolitan vs locally distribution of heterotrophic bacterial OTU isolates

The most abundant and cosmopolitan culturable OTUs and genera, i.e. those that occurred in all or most (around 80%) of the 19 stations studied, and the ones only retrieved locally with a restricted distribution [59] within the samples studied were identified. Seven OTUs were the most abundant ones with more than 50 isolates each (Fig. 4a). From these, only OTU 1, affiliating to *Alteromonas* sp., with 217 isolates, showed a global distribution, appearing in 80% of the stations sampled, and thus, it was truly cosmopolitan. The other six abundant OTUs, affiliating to the genera *Erythrobacter*, *Marinobacter* and *Halomonas*, had a regional distribution having been isolated in more than 25%, but less than 75% of the samples. Moreover, we found 13 less abundant OTUs that also presented a regional distribution and covered different oceanographic regions (Table S2). The remaining OTUs, representing 86.3% of the total OTUs, were isolated locally, that is, in less than 25% of the samples.

**Fig. 4.**
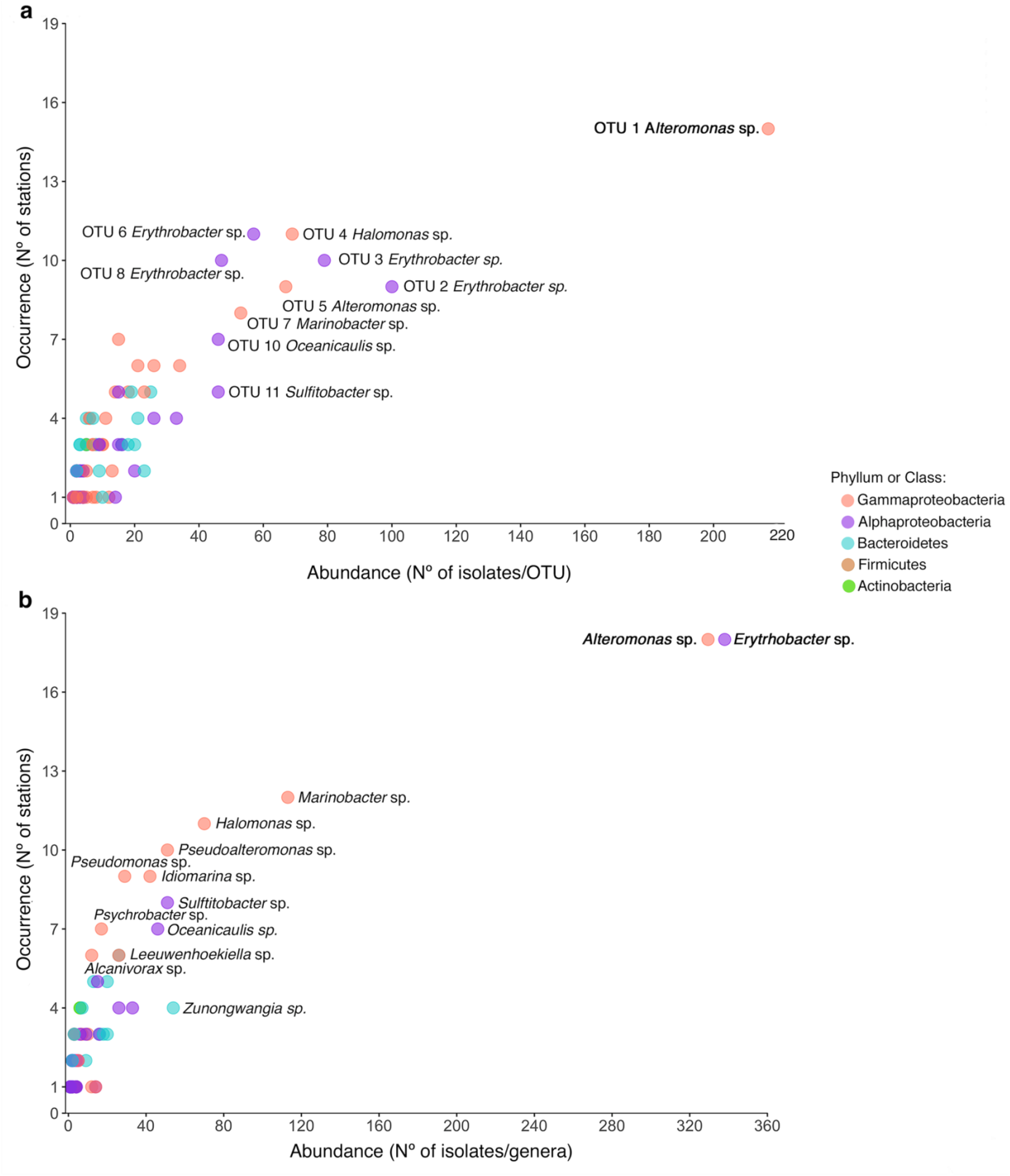
Abundance of the isolates retrieved *versus* their occurrence in the different stations of the study based on: **(a)** OTUs defined at 99% similarity, **(b)** Genera retrieved in the total culture collection. In bold are indicated the most abundant and cosmopolitan OTUs or genera, and in regular type those with a regional distribution. The colour of the dots indicates the taxonomic (phylum or class) affiliation of the OTUs.

*Erythrobacter* and *Alteromonas* were the most abundant and cosmopolitan genera, representing 41.3% of the isolates (338 and 333 isolates respectively), and appearing in 94% of the samples studied (Fig. 4b). Less abundant genera such as *Marinobacter* (113 isolates), *Halomonas* (70 isolates), *Pseudoalteromonas* (51 Isolates), *Idiomarina* (42 isolates), *Pseudomonas* (29 isolate), *Sulfitobacter* (51 Isolates), or *Oceanicaulis* (46 isolates) were present in more than 25% of the samples (Fig. 4b) and covered almost all the oceanographic regions (Table S14). These were, thus, regionally distributed [59]. Some genera such as *Psychrobacter*, *Leeuwenhoekiella* or *Alcanivorax* had lower numbers of isolates, but were recovered in more than 25% of the samples (Fig. 4b). Other genera, in turn, such as *Zunongwangia*, were recovered in less than 25% of the samples but presented 54 isolates (3.5% of the sequences). All the mentioned genera were found in the photic, OMZ, and bathypelagic layers, except for *Oceanicaulis* which could not be retrieved from the bathypelagic samples (Table S14). Finally, the remaining genera represented 20% of the cultures and were recovered only in some of the stations.

### Patterns of genetic distance among culturable heterotrophic marine bacteria

Unweighted UniFrac distances were calculated between pairs of stations and they are expressed with values ranging from 0 (100% similarity) to 1 (0% similarity). Distances between photic-layer stations ranged from 0.31 to 0.79, between OMZ from 0.52 to 0.75, and between bathypelagic samples from 0.48 to 0.8 (Fig. 5a). When we compared the genetic distances between different depths (Fig. 5a), we did not find higher dissimilarities than between those stations coming from the same layer. Besides, ANOVA results indicated that the differences between compared groups were not statistically significant (P-value = 0.08). Furthermore, non-metric multidimensional scaling plot (NMDS, Fig. 5b) of the UniFrac distances did not display a clear separation between samples from different layers. On the other hand, we did not find any correlation between genetic and geographical distances (Fig. 6a). In other words, we did not find higher genetic dissimilarities when the geographic distance between stations increased. The Mantel correlation between the genetic and the environmental matrix considering conservative properties of oceanic waters, such as temperature and salinity, or the chlorophyll *a* (Chl *a*) concentrations, was not significant (r = 0.13, P-value = 0.126). This indicated that environmental parameters did not shape the genetic distances between isolated stations. The distance based redundancy analysis (dbRDA, P-value = 0.058) showed a slight segregation of photic and aphotic samples, but not a clear separation between samples from different oceanographic regions, except for those of the Mediterranean, as seen also in Figure 5b. Temperature, salinity and Chl *a* accounted for a total of 21% of the observed variance with the first two axis explaining 17% (Fig. 6b). Finally, when looking at the bathypelagic samples, the structure of OTUs belonging to individual samples was not evident based on their predominant water masses (Fig. 6c).

**Fig. 5.**
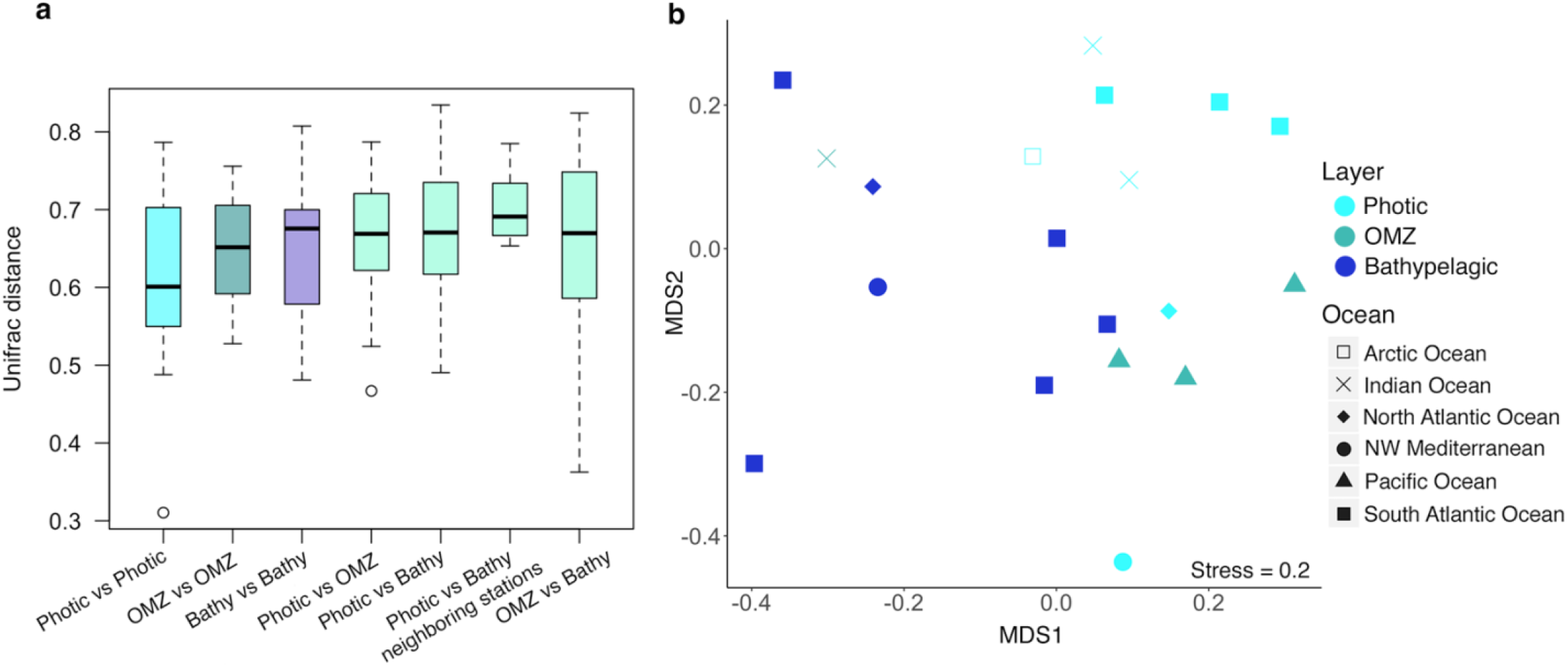
Unweighted UniFrac distances between samples. **(a)** Boxplots showing the distribution of the UniFrac distances between photic-layer, oxygen minimum zone (OMZ), and bathypelagic (Bathy) samples. Same location stands for those photic-layer and bathypelagic stations belonging to neighbouring stations, even though samples were taken in different oceanographic cruises. ANOVA between values of the different groups were not statistically significant (P-value = 0.08). **(b)** Non-metric multidimensional scaling plot indicating the unweighted UniFrac distances between stations. Colour indicates the layer where the stations comes from: light-blue, Photic-layer; turquoise, Oxygen minimum zone (OMZ); and dark blue, Bathypelagic ocean. Shape indicates the oceanographic regions where the station comes from: empty square, Arctic Ocean; cross: Indian Ocean; filled diamond, North Atlantic; filled circle, North Western Mediterranean; filled triangle, Pacific Ocean; and filled square, South Atlantic Ocean.

**Fig. 6.**
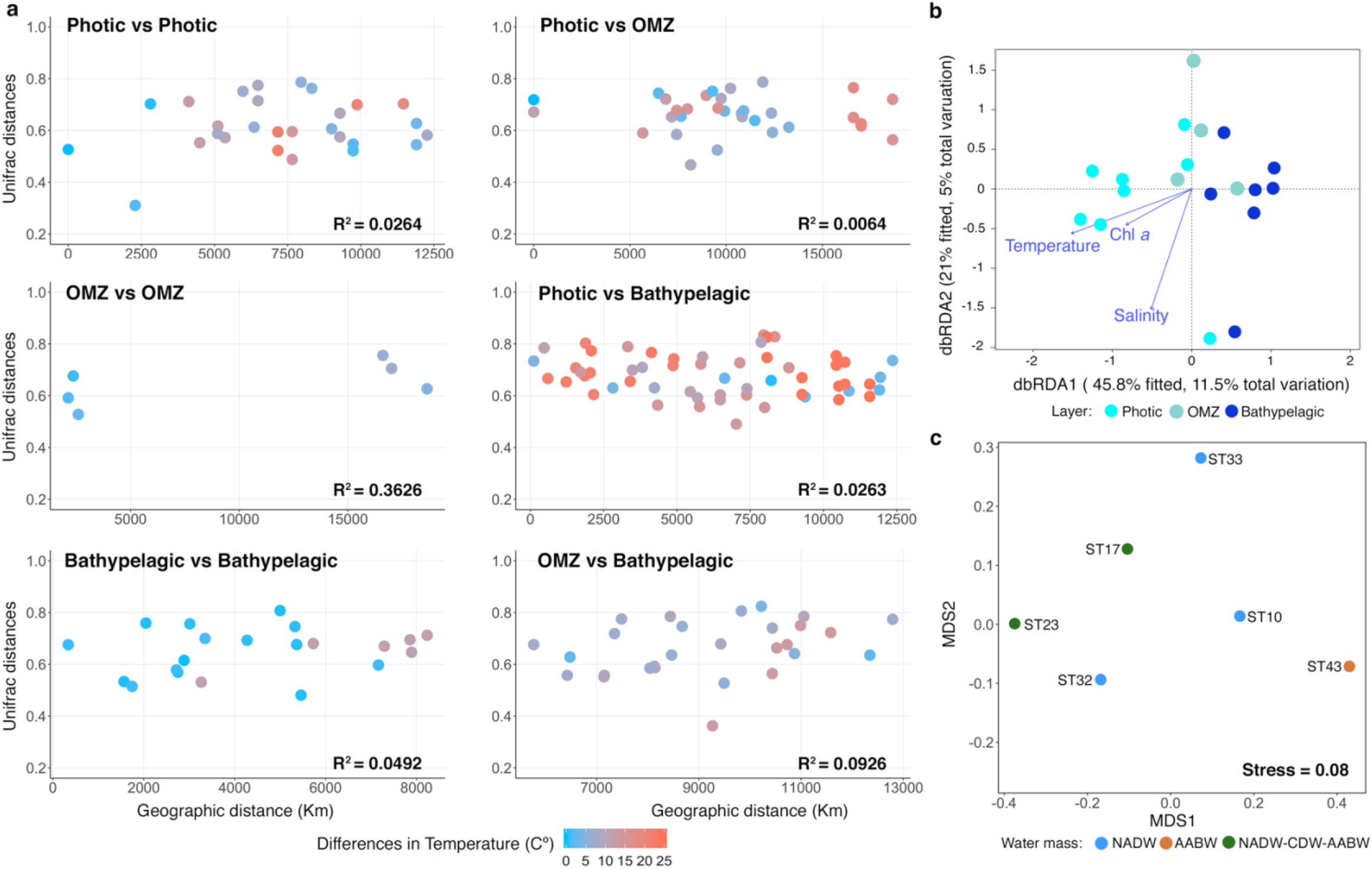
Correlation between UniFrac distances and environmental parameters. **(a)** Graphic representation of the genetic UniFrac comparisons between isolates retrieved from different layers. The color of the points represents differences in the *in situ* temperature between pair of samples. Blue colors indicate less than 10 ºC of difference while orange colors indicate up to 25ºC of difference. R-square values are indicated for each of the graphs. **(b)** Distance-based redundancy analysis of the samples (dots) with three possible explanatory variables (arrows) influencing the genetic distances between isolates of different samples (PERMANOVA p > 0.05; temperature, salinity, and Chlorophyll *a* (Chl *a*) concentration). The ordination was done on the UniFrac distance matrix. Samples are colored by layer: light-blue, Photic-layer; turquoise, Oxygen minimum zone (OMZ); and dark blue, Bathypelagic ocean. **(c)** Non-metric multidimensional scaling plot indicating the unweighted UniFrac distances between only bathypelagic stations. Stress is indicated inside the plot. Colour indicates the predominant bathypelagic water masses of the corresponding stations: NADW, North Atlantic deep water; AABW, Antarctic bottom water, and NADW-CDW-AABW, mix between North Atlantic deep water, Circumpolar deep water and Antarctic bottom water.

Despite the fact that we sampled very distant stations with contrasted environmental conditions and depths (photic vs aphotic), we found that 37% of the isolates (578 out of 1561), affiliating to 10 different genera, were 100% identical at their partial 16S rRNA genes. This percentage increased to 45.7% (374 out of 819 total sequences), affiliating to 13 genera when comparing photic-layer and bathypelagic isolates coming from neighbouring stations, and was even higher, up to 58.9%, when considering all OMZ and bathypelagic samples as aphotic and comparing them to all the photic samples. In all cases, we found *Alteromonas*, *Cobetia*, *Erythrobacter*, *Leeuwenhoekiella, Halomonas*, *Idiomarina, Marinobacter*, and *Mesonia* between the shared genera, indicating a cosmopolitan distribution.

### Novelty of the isolates

The percentages of similarity between the strains and their Closest Cultured Match (CCM) and Closest Environmental Match (CEM) were extracted and compared with the 97% and 99% identity thresholds to explore the possible novelty of our culture collection. The results showed that most of the isolates were similar to previously published cultured bacterial species, but also to environmental sequences obtained using molecular techniques (Fig. 7a). Therefore, most of the isolates were previously known microorganisms. However, we detected three 100% identical strains in their partial 16S rRNA gene that had a percentage of identity, both for CCM and CEM, below the threshold, at around 94%. One of the strains was isolated from surface samples of the North Atlantic Ocean (ISS653), whereas the other two were isolated from two oxygen minimum samples of the Pacific Ocean (ISS1889, ISS2026). Further analyses with the complete 16S rRNA gene of one of those three strains indicated that they could be candidates to be a new specie or even a new genus according to the thresholds proposed by Yarza et al. [60]. The three databases consulted (NCBI, RDP, Silva) showed different BLASTn results (Table S15). Nevertheless, the Living Tree Project (LTP) database, which contains the accepted type species of each genus, showed a 93.5% similarity with *Mesonia mobilis*. The phylogenetic tree constructed (Fig. 7b) also supported its novelty as our isolate had less than 93.8% of similarity with the cultivated reference genomes of the *Mesonia* genus.

**Fig. 7.**
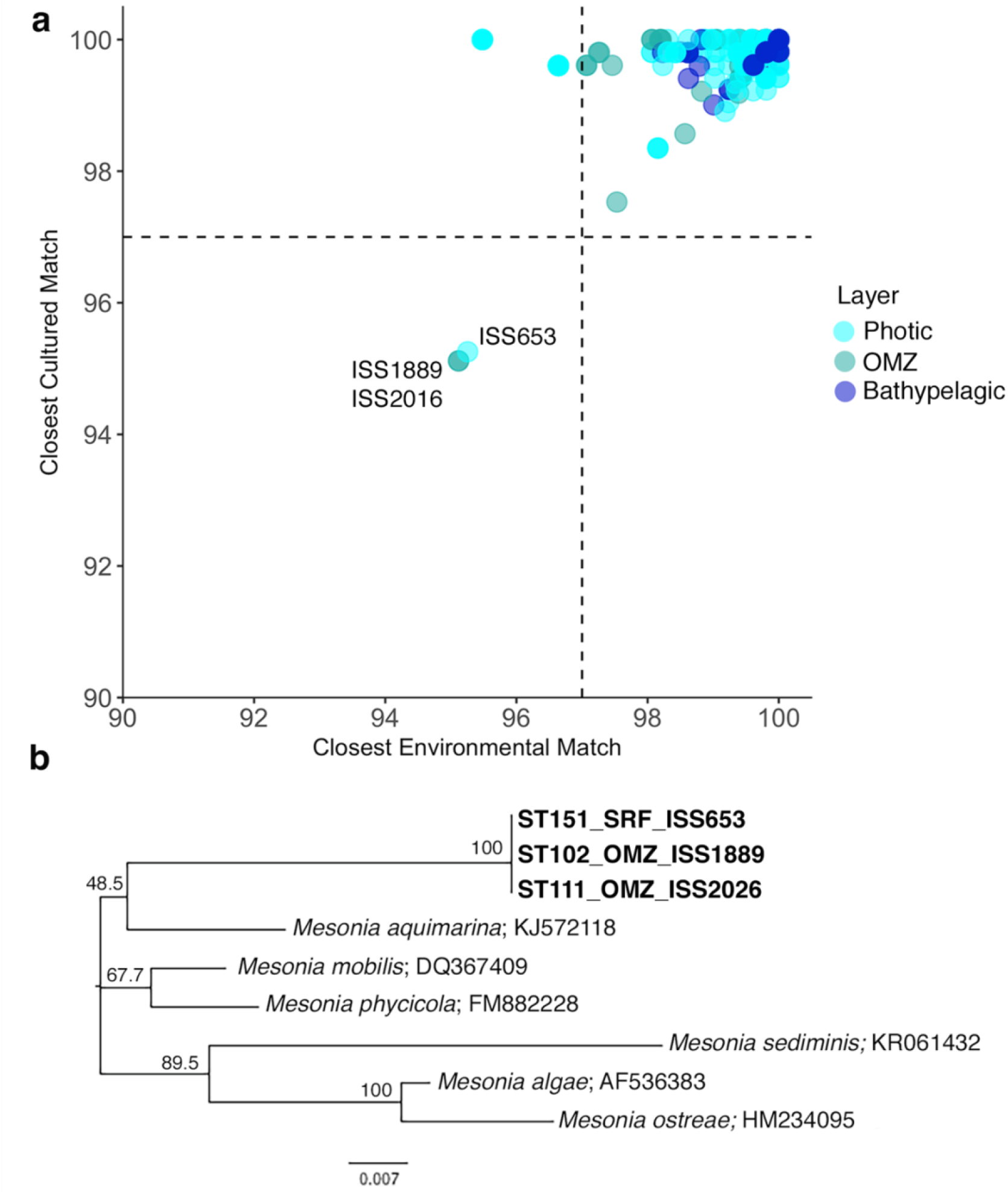
Potentially novel isolates. **(a)** Percentages of similarity between the Closest Cultured Match (CCM) and the Closest Environmental Match of the 16S rRNA gene sequences. Horizontal and vertical lines represent the typical cut-off value of 97% and 99% commonly used for “species” delineation. Colour of the points indicate the layer where the isolates were retrieved from. **(b)** Neighbour Joining tree of the putative *Mesonia* isolates. The numbers in the nodes represent bootstrap percentages > 45%, calculated from 1000 replicates. Putative new isolates are written in bold letters. SRF, surface isolates; OMZ, oxygen minimum zone.

## Discussion

### Richness and abundances of culturable heterotrophic isolates

The rarefaction curves for photic-layer, OMZ, and bathypelagic samples for the isolates dataset did not reach saturation indicating that diversity was higher at all depths. Taking the more restrictive clustering of the isolates sequences (OTUs100), species richness (Chao-1, ACE) increased with depth (from 140 to photic up to 174 at the bathypelagic), and this pattern is coherent with the patterns observed using fragments of 16S rRNA genes directly from metagenomes (16S miTAGs) at global scale [3]. However, Shannon-Weaver and Simpson indices that incorporate both richness and evenness were slightly higher for photic (4) than aphotic layers (OMZ (3.55) and bathypelagic (3.87)). Rank abundance curves from different layers showed that the heterotrophic isolates were composed by a few abundant OTUs with a large number of strains and many rare low abundant OTUs (Fig. 2c-d), which is also consistent with many other previous findings based on environmental samples [1, 8]. For instance, the 7 most abundant OTUs (99%) accounted for 41% of the total sequences and similar proportions were found in each layer. Hence, 30% of the bathypelagic isolates, 47% of the OMZ isolates, and 43% of the photic isolates affiliated with these seven most abundant OTUs.

### Relevance of dispersal along the oceanic water column within heterotrophic culturable bacteria

Small-sized planktonic organisms, such as the prokaryotes, are expected to have a great capacity of dispersion [61–63], more than those of larger planktonic organisms [63, 64]. Indeed, dispersal limitation tends to increase with body size in planktonic communities [62, 63, 65]. Due to the significant overlap between photic, OMZ and bathypelagic isolates, mainly those affiliating to genera *Alteromonas*, *Halomonas*, *Marinobacter* and *Erythrobacter*, our results suggest that these heterotrophic bacteria are well adapted to live in both environments (photic and aphotic), and therefore adapted to different temperatures, light and pressure. Moreover, they probably have versatile metabolisms to respond to different environments and nutrient availability. These characteristics may make these bacteria more prone to be successfully dispersed by the ocean circulation. In addition, genomic comparison between cultured isolates and uncultured genomes retrieved by single amplified genomes (SAGs) from marine environments revealed that the genomes of the cultures had larger sizes, suggesting a predominant copiotrophic lifestyle [66]. One possible explanation supporting the high proportion of identical 16S rRNA gene sequences between isolates of photic and deep-sea layers would be that these bacteria have the capacity to attach and grow on particles in the photic layers and after sinking to the deep ocean, they still retain the capability for further growth. Certainly, a recent study claimed that the particle colonization process that takes places in the photic-layers determines the composition of deeper layers and especially bathypelagic communities, and thus, photic and deep-ocean prokaryotic communities are strongly connected via sinking particles [11]. Moreover, the attachment to particles and its presence in the deep ocean has been described at least for *Alteromonas* sp. [11, 67– 69], *Erythrobacter* sp. and *Halomonas* sp. [11]. The higher metabolic versatility [66] and dormancy capability in the culturable rare bacterial taxa may be crucial to face such long vertical dispersion [70].

Using the partial sequences of the 16S rRNA gene, we did not find significant differences in the genetic distance between isolates that covered a wide latitudinal and geographical distance. Thus, culturable heterotrophic bacteria are not limited by dispersion. Certainly, the timeframes of a few thousand years for complete ocean water turnover [71], could also support the idea of homogenization of microbial populations and the high degree of biogeographical dispersion, as observed for the cosmopolitan genera described. Nevertheless, we also detected taxa present in only a few locations, such as *Vibrio*, isolated only in our survey in the photic North Western Mediterranean Sea, that are probably well adapted to specific niches. In this case, our results are in agreement with the famous sentence **“**everything is everywhere, but the environment selects” [72].

### Extent of cosmopolitan and locally cultured bacteria

As we already mentioned, our results identified *Alteromonas*, *Erythrobacter, Marinobacter* and *Halomonas* as the most abundant cultured OTUs and widely distributed genera in our dataset. We classified the first two as cosmopolitan taxa and the last two as regional taxa, based on the classification proposed in the study of Gregory et al. [59]. These genera have been detected in other culture-dependent and culture-independent studies from a wide variety of marine environments, including coastal, shelf, and open ocean waters [9, 13, 16, 69, 73–75] corroborating their ubiquity.

*Alteromonas* and *Erythrobacter* presented the highest number of isolates. *Alteromonas* is among the most common culturable heterotrophic bacteria living in open marine waters all around the world, as it has been isolated from a wide variety of marine environments [20, 76–79]. In addition, this genus is thought to be one of the most significant contributors of the dissolved organic carbon (DOC) consumption and nutrient mineralization in the upper ocean [80]. *Erythrobacter* strains are aerobic chemoorganotrophs, and some species contain *bacteriochlorophyll a*, responsible for the aerobic anoxygenic phototrophic (AAP) metabolism [75].

Some culturable OTUs had a restricted distribution, appearing only in very low numbers or in only one of the samples. However, some of these, such as the genera *Roseivirga* or *Winogradskyella*, found only in surface waters of the Atlantic Ocean, *Polaribacter*, only retrieved from surface NW Mediterranean waters or *Novosphingobium*, isolated from deep waters of the South Atlantic, had previously been described in other culture-dependent studies from other oceanographic locations [81–85]. Additionally, metadata information retrieved from the Closest Cultured Match and Closest Environmental Match of each one of the strains (Table S3 and S4) also indicated the presence of such restricted genera in other depths and oceanographic regions. Thus, these “local” OTUs are not restricted to the locations where we retrieved them, but the environmental conditions during the sampling and the culturing media used in the study allowed them to be retrieved only from certain samples.

### Novelty in the isolates

There are several well-accepted criteria for the classification of bacteria into species and these include: (i) 70% DNA-DNA hybridization [86], (ii) average nucleotide identity of shared genes (ANI) with a threshold around 94-96% [87], and (iii) 16S rRNA gene sequence identity with a threshold at around 98.7%-99% [46, 60, 88]. In order to further classify bacteria into high taxonomic ranks, Yarza et al. [60] proposed the following taxonomic thresholds for Bacteria and Archaea classification based on 16S rRNA sequence similarity: genera 94.5%, families 86.5%, orders 82.0%, classes 78.5% and phyla 75%. Three strains of our culture collection, 100% similar among them in their 16S rRNA gene, presented less than 95% of similarity in their 16S sequence to any previously described bacterial species and likely represent a novel species or even a new genus, because they are close to the genus threshold. Interestingly, isolate ISS653, was obtained from the North Atlantic waters whereas ISS1889 and ISS2026 were retrieved from oxygen minimum zone areas of the Pacific Ocean, suggesting that this novel species is not locally restricted. The strains possibly belong to the genus *Mesonia*. Members of this taxon are mainly retrieved from a variety of marine environments, sometimes associated with eukaryotic organisms, such as algae [89].

## Conclusions

Culturing remains an important tool in microbial ecology, helping to map the diversity of marine communities, but also to describe the diversity patterns of that fraction of the bacterioplankton community that can be accessed through isolation. Equally to those high-throughput sequencing studies of ribosomal genes targeting the whole marine prokaryotic community, we found that culturable marine heterotrophic bacteria present few abundant taxa and a tail of rare and low abundant OTUs. We detected that half of the total isolates were shared in the three different depth realms, reinforcing the idea of vertical connectivity between the photic and the deep ocean probably through sinking particles. In addition, we identified *Alteromonas* and *Erythrobacter* genera to be cosmopolitan because they were isolated from more than 80% of the studied samples and from all layers. Finally, we did not find any correlation between genetic distances and depth or geographic distance supporting the idea of high dispersal of the marine heterotrophic culturable microorganisms.

## Acknowledgments

We are grateful to Elisabet Laia Sà for her help along the culturing processes and Vanessa Balagué for helping in the laboratory. We thank our fellow scientists and the crew and chief scientists of the different cruise legs for the smooth operation in the ATP, MIFASOL and Malaspina expeditions. We thank the people and sponsors who participated in the *Tara* Oceans Expedition 2009–2013 (http://oceans.taraexpeditions.org) for collecting some of the samples for culturing used in this study. This is *Tara* Oceans contribution paper number XX. The project Malaspina 2010 Expedition (ref. CSD2008–00077) was funded by the Spanish Ministry of Economy and Competitiveness, Science and Innovation through the Consolider-Ingenio programme. Research was mainly funded by grant BIOSENSOMICS “Convocatoria 2015 de ayudas Fundación BBVA a investigadores y creadores culturales” and, MAGGY Plan Nacional I+D+I 2017 (CTM2017-87736-R) to SGA and the King Abdullah University of Science and Technology (KAUST) through baseline funding to C.M. Duarte and a subaward agreement OSR-2014-CC-1973-02 between KAUST and Universitat Autònoma de Barcelona (UAB). Additional funding was provided by projects REMEI (ref. CTM2015-70340-R) from the Spanish Ministry of Economy and Competitiveness, Arctic Tipping Points (ATP, contract #226248) in the FP7 program of the European Union, and DOREMI (CTM2012-34294).

