## Supplementary Information for "Diversity patterns of marine heterotrophic culturable bacteria along vertical and latitudinal gradients"

**This PDF file includes:**

Supplementary Figures

Headings Supplementary Tables

### Supplementary Figures

a

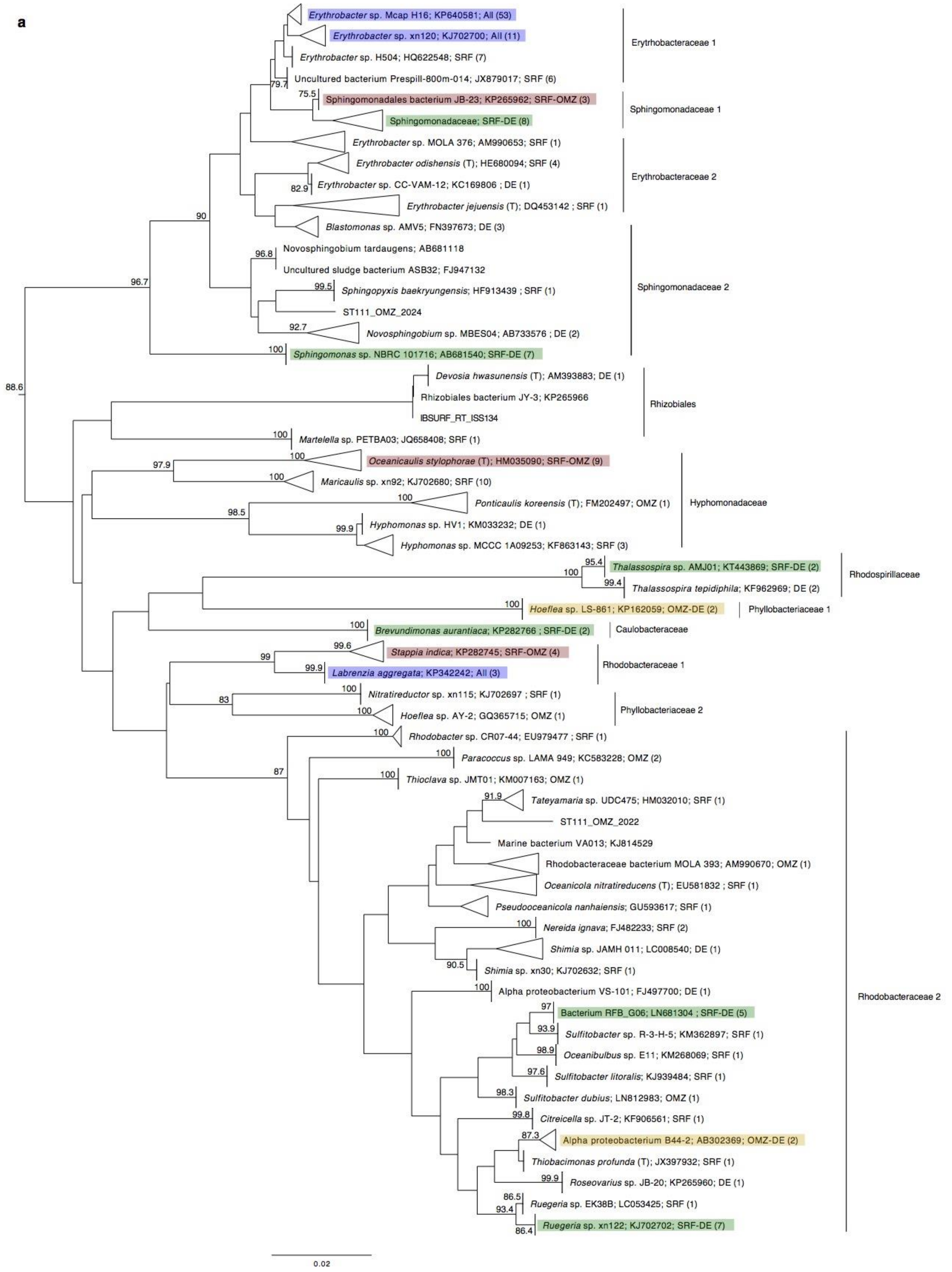

b

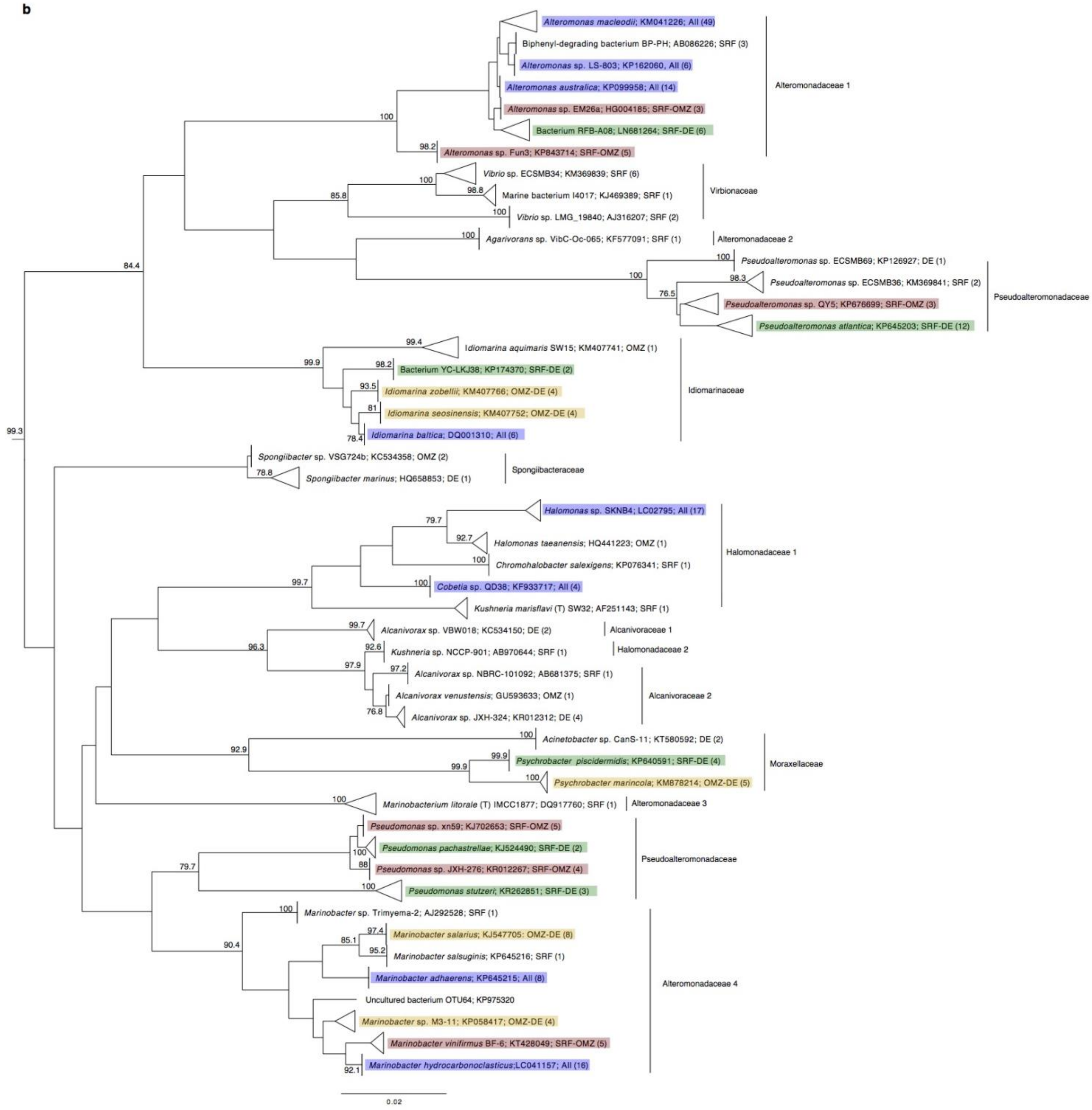

Alteromonas macleodii; KM041226; All (49)
Biphenyl-degrading bacterium BP-PH; AB086226; SRF (3)
Alteromonas sp. LS-803; KP162060, All (6)
Alteromonas australica; KP099958; All (14)
Alteromonas sp. EM26a; HG004185; SRF-OMZ (3)
Bacterium RFB-A08; LN681264; SRF-DE (6)
Alteromonas sp. Fun3; KP843714; SRF-OMZ (5)
Alteromonadaceae 1
Vibrio sp. ECSMB34; KM369839; SRF (6)
Marine bacterium I4017; KJ469389; SRF (1)
Vibrio sp. LMG_19840; AJ316207; SRF (2)
Virbionaceae
Agarivorans sp. VibC-Oc-065; KF577091; SRF (1)
Alteromonadaceae 2
Pseudoalteromonas sp. ECSMB69; KP126927; DE (1)
Pseudoalteromonas sp. ECSMB36; KM369841; SRF (2)
Pseudoalteromonas sp. QY5; KP676699; SRF-OMZ (3)
Pseudoalteromonas atlantica; KP645203; SRF-DE (12)
Pseudoalteromonadaceae
Idiomarina aquimaris SW15; KM407741; OMZ (1)
Bacterium YC-LKJ38; KP174370; SRF-DE (2)
Idiomarina zobellii; KM407766; OMZ-DE (4)
Idiomarina seosinensis; KM407752; OMZ-DE (4)
Idiomarina baltica; DQ001310; All (6)
Idiomarinaceae
Spongiibacter sp. VSG724b; KC534358; OMZ (2)
Spongiibacter marinus; HQ658853; DE (1)
Spongiibacteraceae
Halomonas sp. SKNB4; LC02795; All (17)
Halomonas taeanensis; HQ441223; OMZ (1)
Chromohalobacter salexigens; KP076341; SRF (1)
Cobetia sp. QD38; KF933717; All (4)
Kushneria marisflavi (T) SW32; AF251143; SRF (1)
Halomonadaceae 1
Alcanivorax sp. VBW018; KC534150; DE (2)
Alcanivoraceae 1
Kushneria sp. NCCP-901; AB970644; SRF (1)
Halomonadaceae 2
Alcanivorax sp. NBRC-101092; AB681375; SRF (1)
Alcanivorax venustensis; GU593633; OMZ (1)
Alcanivorax sp. JXH-324; KR012312; DE (4)
Alcanivoraceae 2
Acinetobacter sp. CanS-11; KT580592; DE (2)
Psychrobacter piscidermidis; KP640591; SRF-DE (4)
Psychrobacter marincola; KM878214; OMZ-DE (5)
Moraxellaceae
Marinobacterium litorale (T) IMCC1877; DQ917760; SRF (1)
Alteromonadaceae 3
Pseudomonas sp. xn59; KJ702653; SRF-OMZ (5)
Pseudomonas pachastrellae; KJ524490; SRF-DE (2)
Pseudomonas sp. JXH-276; KR012267; SRF-OMZ (4)
Pseudomonas stutzeri; KR262851; SRF-DE (3)
Pseudoalteromonadaceae
Marinobacter sp. Trimyema-2; AJ292528; SRF (1)
Marinobacter salarius; KJ547705: OMZ-DE (8)
Marinobacter salsuginis; KP645216; SRF (1)
Marinobacter adhaerens; KP645215; All (8)
Uncultured bacterium OTU64; KP975320
Marinobacter sp. M3-11; KP058417; OMZ-DE (4)
Marinobacter vinifirmus BF-6; KT428049; SRF-OMZ (5)
Marinobacter hydrocarbonoclasticus;LC041157; All (16)
Alteromonadaceae 4
100
98.2
100
98.8
100
85.8
100
84.4
100
100
98.3
76.5
99.4
99.9
98.2
93.5
81
78.4
99.3
78.8
79.7
92.7
100
99.7
100
99.7
96.3
92.6
97.9
97.2
76.8
100
92.9
99.9
99.9
100
100
100
79.7
88
100
100
90.4
97.4
85.1
95.2
92.1
0.02

c

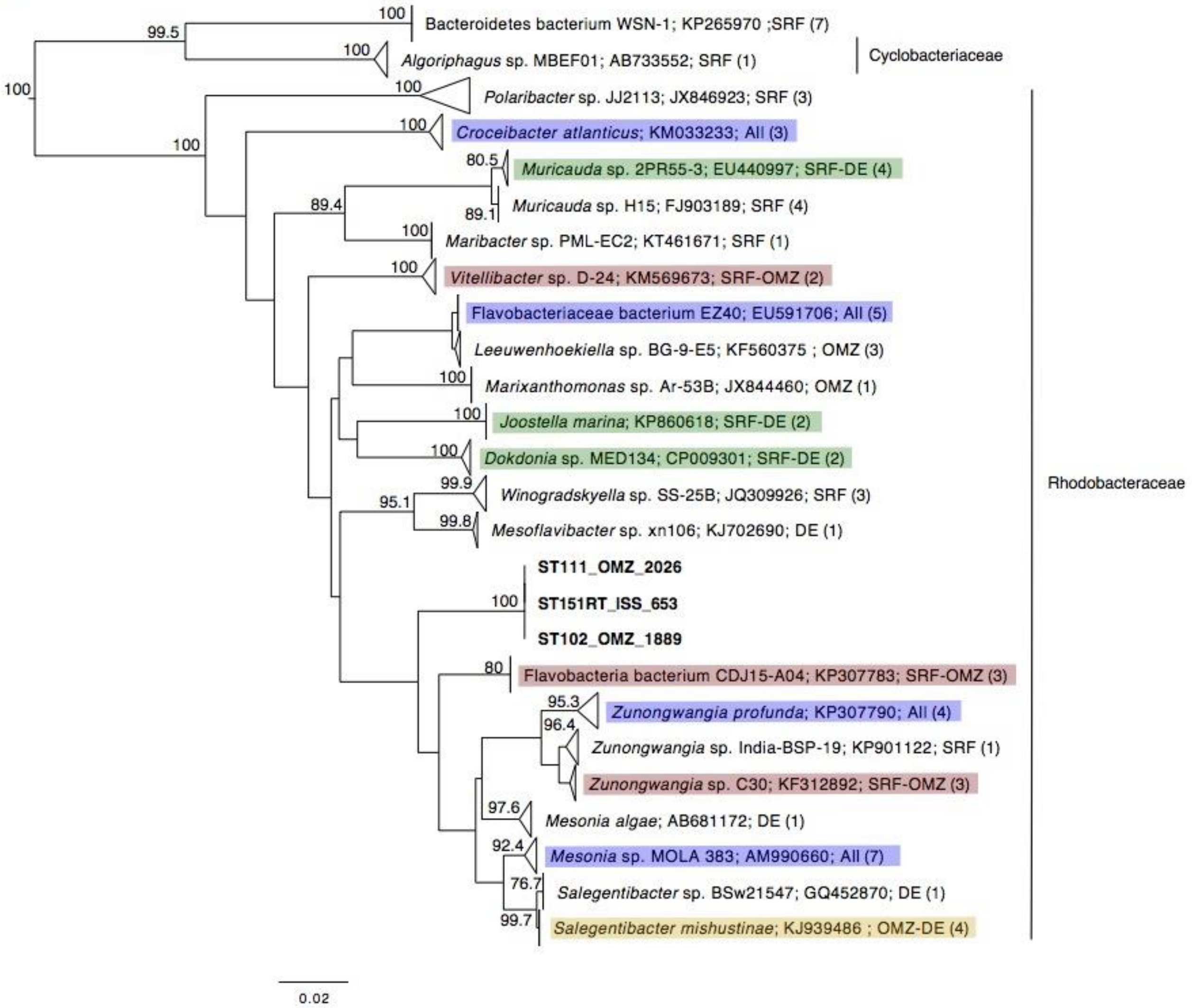

Bacteroidetes bacterium WSN-1; KP265970 ;SRF (7)
Algoriphagus sp. MBEF01; AB733552; SRF (1)
Cyclobacteriaceae
Polaribacter sp. JJ2113; JX846923; SRF (3)
Croceibacter atlanticus; KM033233; All (3)
Muricauda sp. 2PR55-3; EU440997; SRF-DE (4)
Muricauda sp. H15; FJ903189; SRF (4)
Maribacter sp. PML-EC2; KT461671; SRF (1)
Vitellibacter sp. D-24; KM569673; SRF-OMZ (2)
Flavobacteriaceae bacterium EZ40; EU591706; All (5)
Leeuwenhoekiella sp. BG-9-E5; KF560375 ; OMZ (3)
Marixanthomonas sp. Ar-53B; JX844460; OMZ (1)
Joostella marina; KP860618; SRF-DE (2)
Dokdonia sp. MED134; CP009301; SRF-DE (2)
Winogradskyella sp. SS-25B; JQ309926; SRF (3)
Mesoflavibacter sp. xn106; KJ702690; DE (1)
ST111_OMZ_2026
ST151RT_ISS_653
ST102_OMZ_1889
Flavobacteria bacterium CDJ15-A04; KP307783; SRF-OMZ (3)
Zunongwangia profunda; KP307790; All (4)
Zunongwangia sp. India-BSP-19; KP901122; SRF (1)
Zunongwangia sp. C30; KF312892; SRF-OMZ (3)
Mesonia algae; AB681172; DE (1)
Mesonia sp. MOLA 383; AM990660; All (7)
Salegentibacter sp. BSw21547; GQ452870; DE (1)
Salegentibacter mishustinae; KJ939486 ; OMZ-DE (4)
Rhodobacteraceae
100
99.5
100
100
100
100
80.5
89.4
89.1
100
100
100
100
100
99.9
95.1
99.8
100
80
95.3
96.4
97.6
92.4
76.7
99.7
0.02

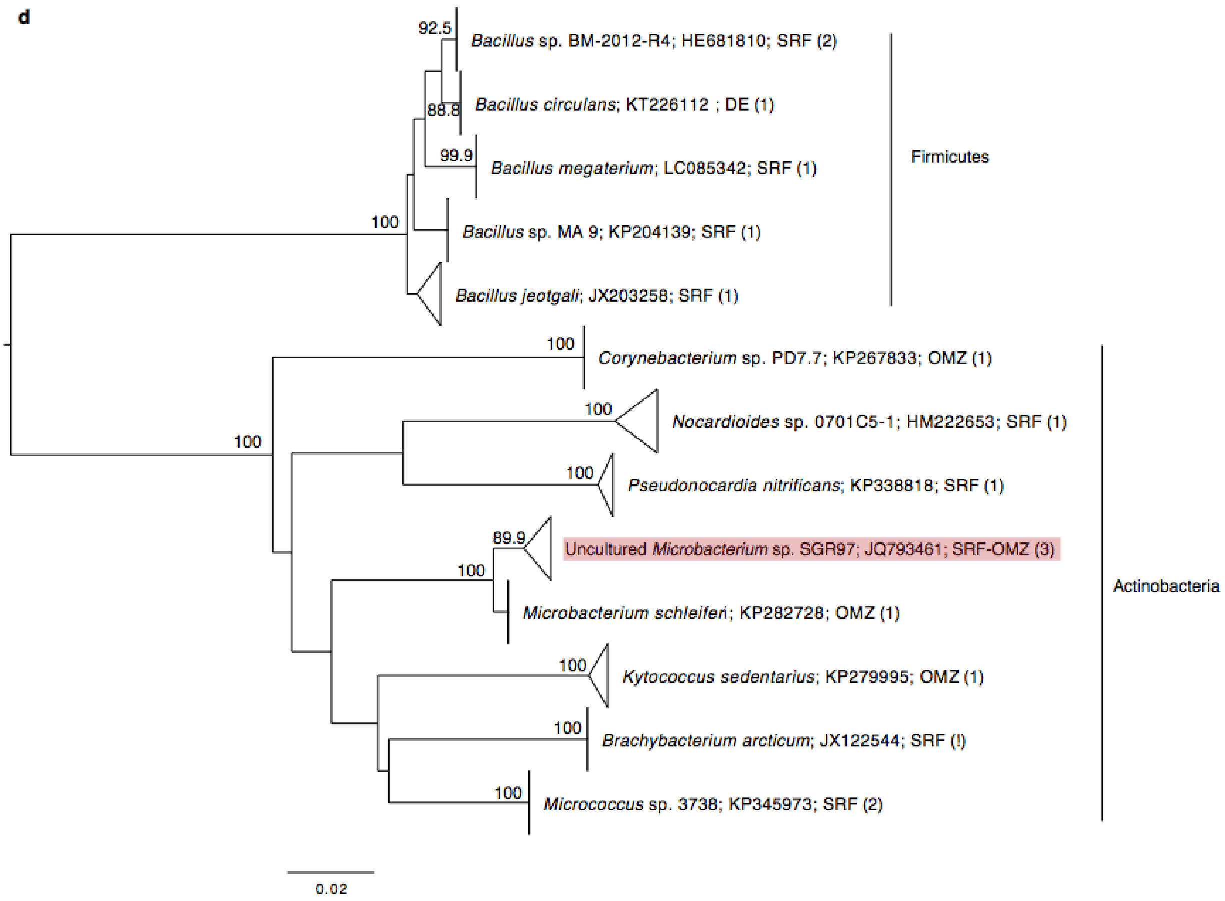


**Fig. S1** Phylogenetic relationships between photic-layer, oxygen minimum zone (OMZ) and bathypelagic isolates**.** Neighbour Joining trees of the 16S rRNA gene sequences of the reduced pool of sequences, including the non-redundant 516 isolates plus their closest cultured match (CCM) and closest uncultured or environmental match (CEM). The numbers in nodes represents bootstrap percentages > 75, calculated from 1000 replicates. The number of isolates from each specific cluster is indicated in brackets. Isolates in bold (only in Bacteroidetes tree) show the putative novel genera isolated in this study. Blue rectangles indicate group of isolates retrieved from all depths; in red; a mix between photic-layer and OMZ; in green, a mix from photic-layer and bathypelagic isolates; and in yellow, a mix of OMZ and bathypelagic. DE, bathypelagic isolates; SRF, photic-layer isolates; OMZ, oxygen minimum zone isolates. The vertical lines indicate the family name or order of some groups of isolates in each tree. **(a)** *Alphaproteobacteria*; **(b)** *Gammaproteobacteria*; **(c)** *Bacteroidetes*; **(d)** Gram positive bacteria.

**Headings Supplementary Tables**

**Table S1.** Culture media and incubation conditions used for each seawater sample. Positive signs indicate which media where used. RT: room temperature.

**Table S2.** Non-subsampled OTU-abundance table per depth defined at 99% sequence similarity. SRF: photic-layer; DE: bathypelagic; OMZ: oxygen minimum zone.

**Table S3.** Metadata information of the Closest Cultured Match (CCM) obtained after BLASTn analysis of the isolates against a subset of the RDP database including only sequences from previously published cultured bacteria. SRF: photic-layer; DE: bathypelagic; OMZ: oxygen minimum zone.

**Table S4**. Metadata information of the Closest Environmental Match (CEM) obtained after BLASTn analysis of the isolates against a subset of the RDP database including only sequences from previously published uncultured bacteria. SRF: photic-layer; DE: bathypelagic; OMZ: oxygen minimum zone.

**Table S5.** Comparisons between the number of OTUs and the percentage of shared sequences between photic-layer and bathypelagic samples from near locations in non-subsampled and subsampled OTU tables.

**Table S6.** Subsampled OTU-abundance table including only the photic-layer and bathypelagic samples from near locations. OTUs obtained after clustering at 99% sequence similarity. SRF: photic-layer isolates; DE: bathypelagic isolates.

**Table S7**. Table indicating the number of total isolates affiliating to each genus found in the photic and bathypelagic samples from close locations. Results obtained after grouping all the OTUs from the subsampled OTU-abundance table (99% clustering) affiliating with the same genus. SRF: photic-layer isolates; DE: bathypelagic isolates.

**Table S8.** Fisher tests results indicating overrepresentation of certain genera in the photic or bathypelagic layer. SRF: photic-layer isolates; DE: bathypelagic isolates.

**Table S9.** Comparisons between the number of OTUs and the percentage of shared sequences between photic-layer, oxygen minimum zone (OMZ) and bathypelagic samples in non-subsampled and subsampled OTU tables.

**Table S10.** Comparisons of the richness and diversity indexes estimated using the isolates OTU tables per layer defined at 100% and 99% sequence similarity with and without subsampling down to the lowest isolated depth. No R., stands for non-rarefied or non-subsampled OTU table, and R., stands for rarefied or subsampled OTU table.

**Table S11.** Subsampled OTU-abundance table including all isolates from the photic, the OMZ and the bathypelagic. OTUs obtained with clustering at 99% sequence similarity. SRF: photic-layer; DE: bathypelagic; OMZ: oxygen minimum zone.

**Table S12.** Table indicating the number of total isolates affiliating to each genus found in the photic, OMZ, and bathypelagic samples. Results obtained after grouping all the OTUs from the subsampled OTU-abundance table (99% clustering) affiliating with the same genus. SRF: photic-layer isolates; OMZ: oxygen minimum zone; DE: bathypelagic isolates.

**Table S13.** Summary of the Fisher test analysis indicating which genera had been found overrepresented in the photic, OMZ or bathypelagic layer.

**Table S14.** Table indicating the number of total isolates affiliating to each genus found in the photic, OMZ, and bathypelagic samples. Results obtained after grouping all the OTUs from the non-subsampled OTU-abundance table (99% clustering) affiliating with the same genus. SRF: photic-layer isolates; OMZ: oxygen minimum zone; DE: bathypelagic isolates.

**Table S15.** New potential isolates best hits. Closest Cultured Match (CCM) and Closest Environmental/Uncultured Match (CEM) of the ISS653, ISS1889 and ISS2026 isolates BLASTn results against the NCBI, RDP 11 and SILVA LTP databases. Accession number and percentage of similarity are indicated together with the best hit.
